# Epithelial polarity drives tissue tension in planar, free standing cell monolayers

**DOI:** 10.1101/2025.03.06.641857

**Authors:** Amaury Perez-Tirado, Ulla Unkelbach, Andreas Janshoff

## Abstract

Polar epithelial cells form thin but resilient sheets that resist mechanical in-plane stress by relying on strong conformal contacts with each mediated by dedicated cell-cell connections connected to the viscoelastic cortex. In this study, we investigate the mechanical response of free-standing cell monolayers to central indentation as a function of orientation using a colloidal probe. We determine tissue tension by treating the deformed tissue as a minimal surface area. Our findings reveal that the cortex tension of the basal side governs the purely elastic response to in-plane extension, while the apical side of the cells is soft and dissipative giving rise to a hysteresis at low indentation depth. At larger indentation depth, the apico-basal polarity is no longer relevant as the cells are apically compressed and the response is driven by the elastic in-plane response of the basal side of the tissue. These results are particularly significant for lumen-forming epithelial cells, which experience substantial compressive forces especially apically due to elevated Laplace pressure.

## INTRODUCTION

Epithelial monolayers, whether as lining body cavities or 3D structures, serve as physical and chemical barriers, managing the permeability between the internal and external environments. These tissue sheets’ function is often linked to their remarkable mechanical resilience.^1–3^ To achieve optimal tissue resilience, the cells need to be mechanically interconnected to spread stresses across the entire tissue. This is accomplished through flexible, viscoelastic cell-cell junctions such as desmosomes, adherens junctions, and tight junctions, which link elements of the cytoskeleton to form structures that surpass the individual cell scale. These laterally interconnected, coherent sheets display apico-basal polarity, which is pivotal not only for fulfilling diverse cellular functions, from barrier to the positioning of the adhesion molecules.^4–7^ The development of a polarized epithelial phenotype entails a significant reorganization of the cytoskeleton, polarized membrane trafficking, the creation and maturation of cell junctions, signaling pathways, and the establishment of cortical phospholipid asymmetry.^8–10^

In the scenario of 3D epithelial structures, such as renal tubes, the apical cell surface is embellished with microvilli and is oriented towards the lumen of a tubule. In contrast, the basolateral cell surface is firmly anchored to the extracellular matrix (ECM) and forms lateral connections with neighboring cells of the monolayer. This polarization is achieved by a distinctive protein and lipid composition of the two surface domains mirroring the functional disparities existing between them in the epithelium.^11–13^ Despite the fact that the molecular organization of basal and apical membranes is actively maintained, our knowledge regarding mechanical differences between the apical and basal cortices remains sparse.^14^ Recent findings by Fischer-Friedrich and colleagues reveal that the basement membrane exhibits elastic sheet-like behavior. Their study highlights the crucial roles of both the basement membrane and the actomyosin network as key structural elements in generating basal tension.^15^

Furthermore, polarization is essential for the existence of complex multicellular organisms with diverse cell types. Mutations or pathogens affecting the cytoskeleton, polarized structure, or junctions can increase epithelial tissue fragility, leading to developmental defects, inflammation, and cancer.^4, 13, 16^ Previously, Fouchard *et al*. showed that epithelial monolayers curl basally due to asymmetric contractility generating high spontaneous curvature of the sheet.^17^ Blonski *et al*. designed a microfluidic system that initiates epithelial folding, simulating the typical deformations observed during embryogenesis. They discovered that the direction of folding, whether apical or basal, has an impact on the propagation of folding-induced calcium waves.^18^

Here, we examine the mechanical response of circular free-standing cell monolayers to central indentation both apically and basally with a colloidal probe using an atomic force microscope. Tissue tension is obtained from energy minimization rendering the shape of the deformed tissue as a minimal surface area. We find that the response of the cells to deformation is purely elastic and governed by the basal side of the cells. Conversely, the apical side is soft and dissipative but after compression of the apical caps, the basal side dominates the force response. This finding has implications especially for lumen-forming epithelial cells, which are exposed apically to substantial compressive forces due to elevated Laplace pressure.

## Results

### Generation of Free-Standing Cellular Monolayers

Epithelial cells’ apical and basal domains have diverse functions and exhibit distinct mechanical properties. To measure the force response in these domains individually, we cultured the cells to form self-supporting monolayers. We initially devised a small-scale testing setup for creating these free-standing MDCK (Madin-Darby canine kidney) II monolayers, using the method detailed by Harris *et al*.,^4^ which involves cultivating the monolayer on a removable collagen matrix spanned between the two rods (Fig. SI1). Subsequent removal of the collagen matrix is achieved by collagenase treatment (Fig 1A). By using epifluorescence microscopy, we confirm that the monolayer preserves its apico-basal polarity (Fig 1B/C), rendering it suitable to mechanically target the two different sides of the monolayer. However, this setup turned out to be unsuitable for mechanical measurements using colloidal probe microscopy with conventional atomic force microscopes and also mathematically more difficult to describe due to lack of symmetry and complicated boundary conditions (*vide infra*). Hence, we established an alternative method to create circular free-standing cell monolayers accessible by an atomic force microscope from both apical and basal side referred to as the grid mount. The device consists of a circular glass ring to which the cell monolayer is attached and a metal grid to initially host the collagen and thereby support the composite (Fig 1D and Fig SI2). After cells are grown to confluency, the device permits us to completely remove the collagen matrix keeping the cells intact and providing sufficient access of the colloidal probe and cantilever to both apical and basal domain of the cells (Fig 1D/E).

**Figure 1.**
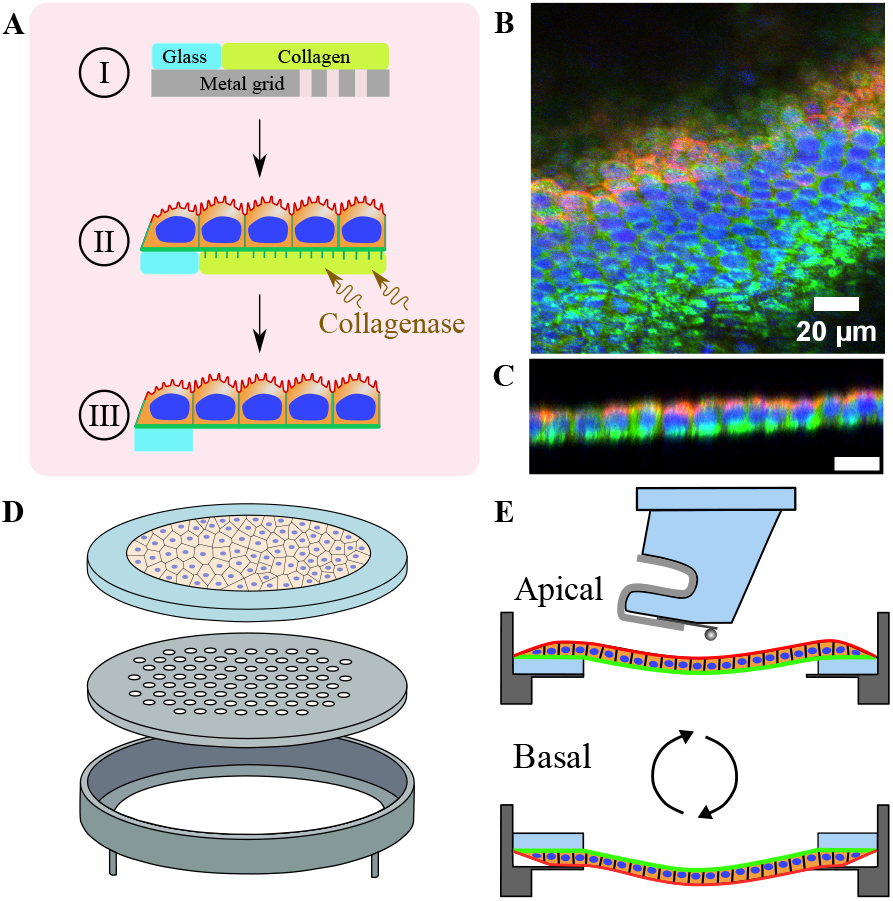
Preparation of free-standing cell monolayers on a grid device. A) Placement of the collagen at the level of the glass ring (I) and subsequent cell seeding (II). Treatment with collagenase to remove the matrix, resulting in a cell monolayer anchored at the edges of the glass without extra support (III). B) Confocal image of an immunostained MDCK II monolayer after removal of the collagen matrix. Anti-podocalyxin is used to label the apical domain (red), while anti-laminin highlights the basal domain (green). DAPI is used to stain the nuclei (blue). Note that the surface is uneven generating a gradient of the different layers of the cells. C) Orthogonal view of the immunostained MDCK II monolayer. Scale bar: 20 μm. D) Schematic illustration of the grid device, consisting of a glass ring where the cells attach basally, a metal grid to initially hold the collagen and the support of the structure. E) Orthogonal view of the device (cross section), showing how turning the device gives access to both sides, apical and basal.

### Mechanical Properties of free-standing Cell Monolayer

Two types of mechanical experiments, indentation-relaxation and indentation-retraction, are conducted with the colloidal probe microscope using spherical tips (20 Âţm in diameter). In indentation-relaxation experiments, the cell monolayer is indented at a constant speed of 2 Âţm/s, followed by a 10-second time period allowing the force to relax at a constant distance. The cycle concludes with the cantilever retracting from the cells at the same speed. Indentation-retraction experiments omit the force relaxation step and involve immediate retraction of the z-piezo. Fig. 2A shows indentation-relaxation curves as force versus time data for cell monolayers after collagen removal deformed apically (left, red) and basally (right, green), respectively. Different maximal force were used to illustrate that the force response to indentation is largely linear even up to 200 nN. Force relaxation after reaching the maximal force (setpoint) indicates viscoelastic behavior in the case of apically loaded monolayers, while basally deformed tissue responds purely elastic, i.e., no change of the force at constant indentation depth.

**Figure 2.**
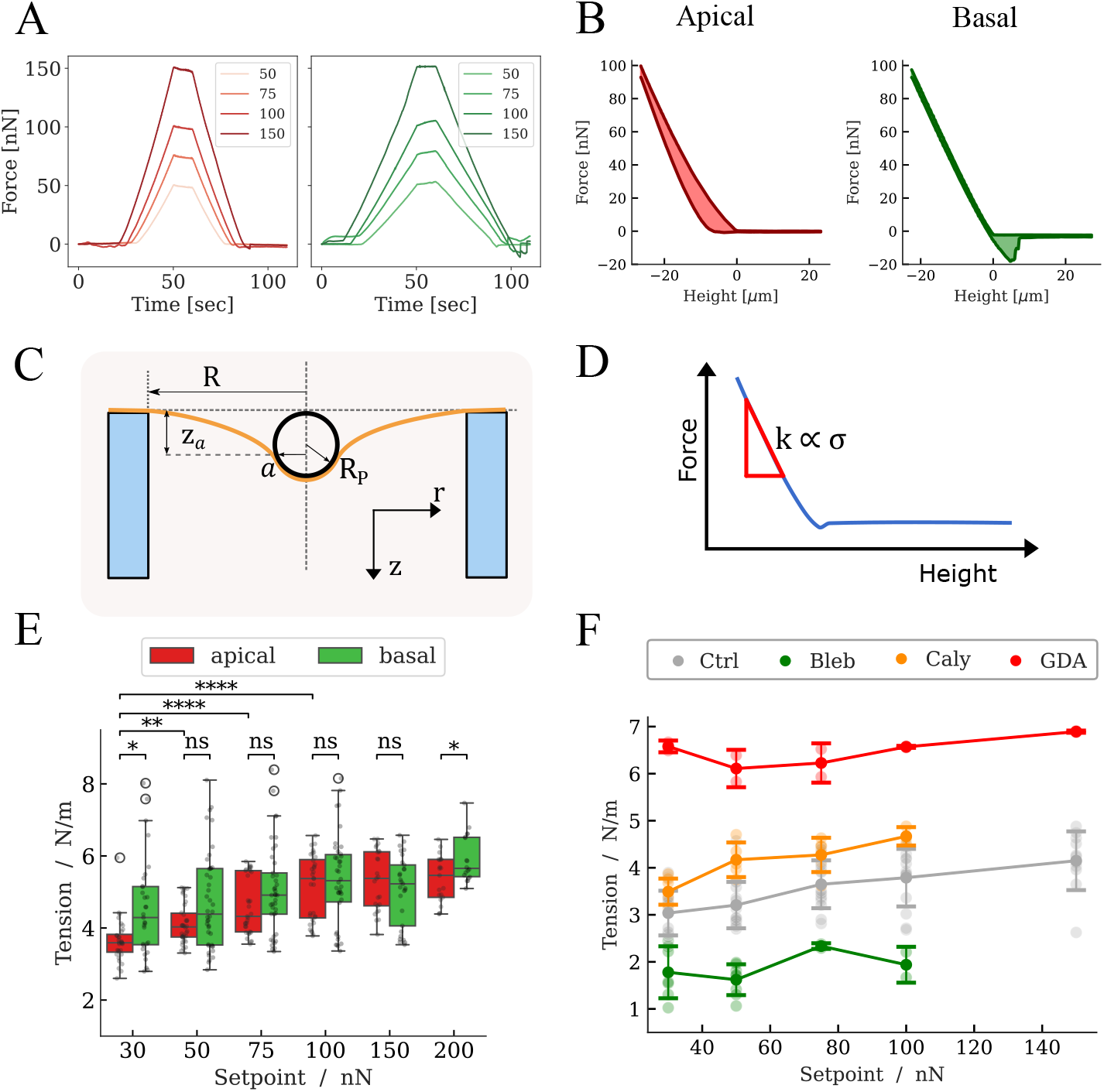
Indentation response of the monolayer. A) Indentation-relaxation experiments (force-time curves) of the apical (left) and basal (right) side of a free-standing cell monolayer loaded up to different setpoints. The response of the basal domain is purely elastic, while the apical side shows slow force relaxation. B) Indentation-retraction experiments (force-height curves) of the apical (left) and basal (right) side of a cell monolayer up to a setpoint of 100 nN. C) Model of the indentation experiment. The yellow line represents the cell monolayer adopting a catenoid shape and the black circle denotes the indenter. *R* is the radius of the aperture covered by the monolayer (inner circle of the glass ring), *z*_*a*_ is the indentation depth at *r* = *a, R*_p_ the radius of the indenter, and *a* is the contact radius of the cell monolayer with the indenter. D) The force versus indentation (height) plot illustrates how the tissue tension is obtained as the slope of the curve. E) Tissue tension is derived from force curves with varying maximal indentation forces. At higher maximal forces (setpoints), cell monolayers loaded apically and basally exhibit identical tissue tension. F) Tissue tension as a function of drug administration with a following probing from the apical side of the cell monolayer. Crtl: MDCK II cell monolayer in the absence of drugs (control); Bleb: blebbistatin (50 μM, stalling myosin II motors); Caly: calyculin A (20 nM, inhibiting phosphatase); GDA: glutardialdehyde (fixation).

Fig. 2B shows two typical indentation-retraction curves (force-height representation) probing a free-standing cell monolayer in the absence of collagen up to approximate 25 Âţm indentation depth (setpoint: 100 nN). On the left side, the cell monolayer is first deformed apically (red) and on the right side the same monolayer is loaded basally (green). Cells deformed basally respond elastically and do not produce a noticeable hysteresis between approach and retraction, while apically loaded cells are initially softer and dissipate energy at these low indentation depths. This is also reflected in the lack of force relaxation found in indentation-relaxation experiments (Fig. 2A) at constant distance. At larger strain the hysteresis between approach and retraction eventually vanishes (Fig. SI5) and the cells’ response becomes stiffer showing a linear dependence of force on indentation. The slope is almost identical to the one found for basally loaded monolayers (Fig. 2A, SI5). Resorting to a continuum view, we treat the cell monolayer as a continuous, homogeneous sheet with a thickness that is negligible compared to the aperture size. We need to consider three fundamental contributions to the free energy that are relevant, i) bending of the thin sheet, ii) in-plane stretching and iii) pre-stress or tissue tension. Bending of the sheet can be safely neglected as a major contribution to the force response because the curvature change is extremely small in these experiments as the radius of the aperture is very large (millimeter range compared to micrometer thickness of the monolayer). In-plane stretching can also be discarded since the probed area *A*_0_ = *πR*^2^ of the aperture is very large compared to the indentation depth (Fig. 2C) leaving the relative area dilatation *α* = Δ*A*/*A*_0_ extremely small. Although the area compressibility modulus is presumably in the range of 0.1 N/m, the small area change still justifies ignoring the stretching of the monolayer during deformation as a contribution to the overall free energy. Hence, we identify tissue tension or pre-stress as the main source of the force response. If a constant tissue tension is responsible for the observed force response, free energy minimization is identical to finding the minimal free surface area (see materials and methods).^19^ Minimization of the unsupported monolayer area leads to a shape representing a catenoid formed between two rings (Fig. 2C), one ring provided by the aperture and the other one the formed by contact of the cells with the spherical indenter. Consequently, the force response during the approach ramp *f*_app_ depends approximately linearly on the indentation depth *z*_max_ (see Fig. 2C/D).

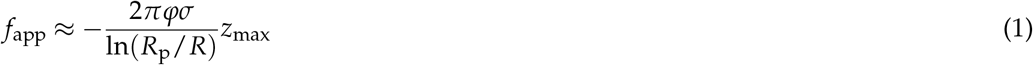

*R* is the radius of the aperture, *R*_p_ the radius of the spherical probe attached to the cantilever, and *σ* the tissue tension. *φ* is an empirical correction factor accounting for the deviation from the full theory (see material and methods). Equation (1) holds if the contact radius equals the indenter radius *a* = *R*_p_, which is an approximation valid for large enough indentation depth *z*_max_ (see materials and methods).

Fig. 2A and Fig. SI5 confirm the linear response up to a maximal force of 200 nN for both apically and basally deformed epithelial monolayers. The tension of free-standing MDCK II monolayers obtained from the slope is in between 3-4 mN/m in good accordance with data from cysts deformation (Fig. 2E).^20^

In order to exclude that the force curves are affected by residual collagen gel, we first examined the mechanical properties of neat collagen membranes, cell monolayers on still existing collagen support and cell monolayers in the absence of collagen after collagenase treatment. Exemplary indentation-relaxation (force-time curves) and indentation-retraction experiments of the collagen matrix and a cell layer on top of the collagen matrix for different setpoints are displayed in Fig. SI3/4. We found that the collagen gel alone shows a typical viscoelastic response with a pronounced hysteresis due to energy dissipation during deformation. This is in line with previous works showing this in bulk rheology experiments.^21^ In the presence of cells, the force response of the composite changes substantially, showing stiffening of the sample by a factor of 2 compared to neat collagen, and concomitantly the dissipated energy is slightly increased for the combined layer. In contrast, in the absence of collagen due to collagenase treatment, no dissipation occurs if probed basally (Fig. 3A/B). Apically, an initial dissipation is always visible attributed to the deformation of the cells themselves (*vide infra*). At higher strains (indentation depths) the dissipation vanishes also in apically loaded tissue, in contrast to the collagen response, where the hysteresis is still increasing (see Fig. SI3-5). In summary, collagen removal with collagenase is sufficient to suppress any mechanical response from residual collagen.

**Figure 3.**
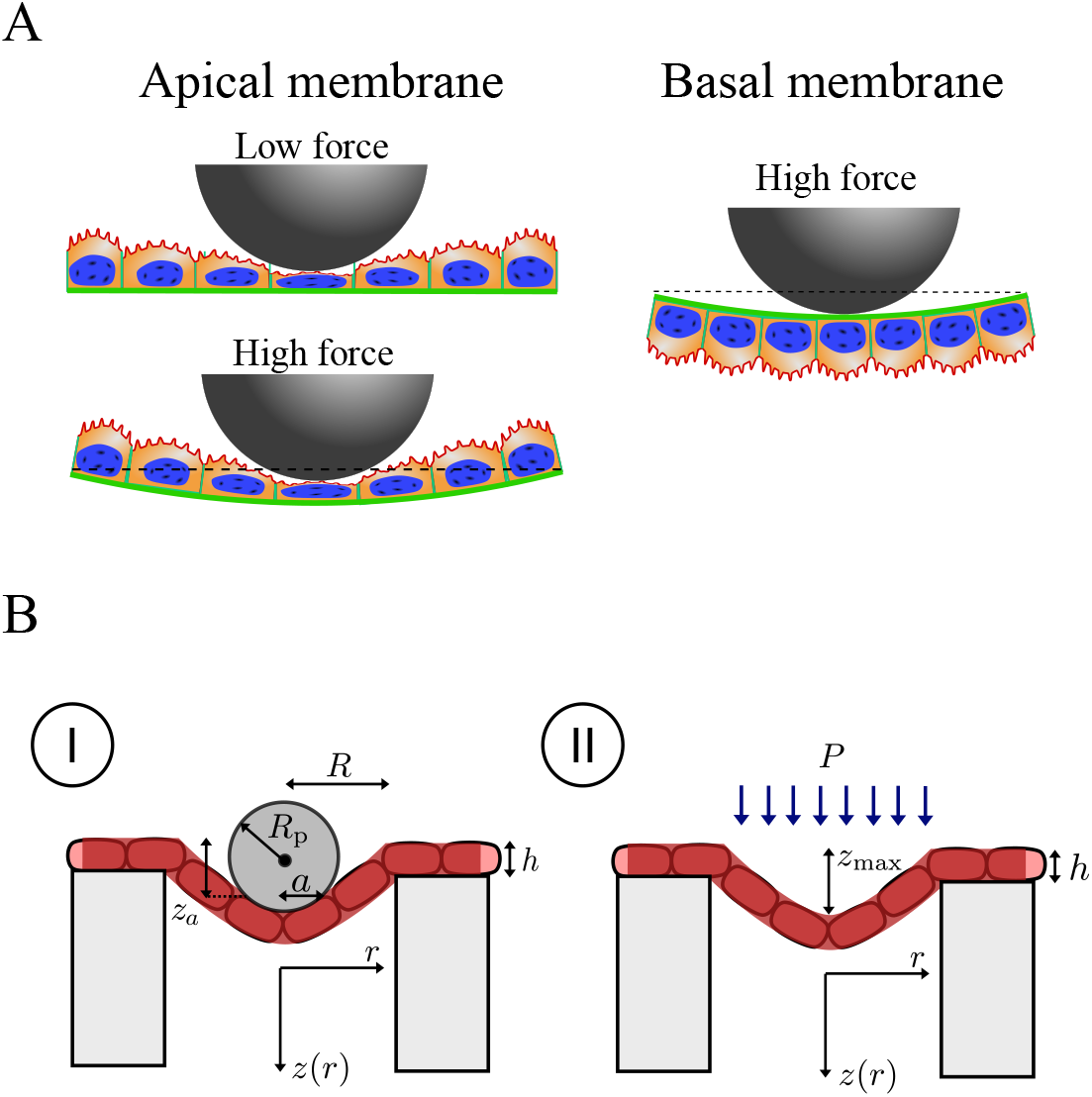
A) Schematic of the mechanical response of the monolayer, the apical side absorbs the deformation at small indentations and gradually start deforming in larger indentations. The basal side is rigid and deforms instantaneously. B) Parametrization of the free-standing cell monolayer subject to indentation with a spherical indenter (I) and gravity (II).

Furthermore, we found that the cell monolayer exhibits a certain amount of slack due to its increased density compared to the culture medium. This slack is estimated both by means of imaging as well as from assuming homogeneous pressure acting on the free-standing monolayer (Fig. 3B (II)) using the tension values obtained from force curves. Essentially, gravity is pre-deforming the cell-monolayer by approximately only 80-100 Âţm. Compared to the aperture size in the millimeter range this can be neglected as the possible source of error for determining tissue tension.

Fig. 2E shows how the tension depends on the polarity and the maximal applied force (setpoint). Here is visible that probing the apical side atenuates the force response as the apical caps are easily deformable giving rise to energy dissipation as shown in Fig. 2B (left). We attribute this energy dissipation to the initial compression of the soft apical caps of the cell bodies.^22^ After apical compression, the response becomes elastic originating from the tissue tension of the entire monolayer. Eventually, at higher setpoints (larger indentation depth) the response from apically and basally deformed tissue sheets converge (Fig. 2E). Hence, at setpoints larger than 100 nN a constant tissue tension is found that does not depend on the orientation of the cell monolayer. This means that eventually at larger forces, the stiffer basal side governs the force response as illustrated in Fig. 2E and Fig. 3A. We suggest that apically deformed cell monolayers dissipate energy through local compression in the contact zone with the spherical indenter, resulting in a softer and initial nonlinear response (Fig. 2B (left)). As the indentation depth increases and the compression of the apical caps ceases, the stiffer and purely elastic basal side becomes dominant (Fig. SI5).

Tissue tension can be modulated by the actomyosin contractility of the cells. While blebbistatin (para-aminoblebbistatin) stalls myosin motors,^23^ calyculin A, a phosphatase inhibitor, increases actomyosin contractility.^24^ Charras and coworkers showed that intrinsic tension increase can lead to local rupture events. We observe a similar behavior and can prove that the overall tissue tension increases with addition of calyculin (Fig. 2F).^4, 6, 7^ Conversely, tension is released by administration of blebbistatin confirming that subcellular perturbations can change the properties of tissue on much larger length scales. Cross-linking the proteins using glutardialdehyde leads to the expected stiffening of the tissue (Fig. 2F). Notably, we still do not probe the bending modulus of the sheet but the tension, which means that motor proteins are captured in the rigor state keeping tension at the highest possible level.

## Discussion

Epithelial cells are the predominant cell type in all animals. They typically form sheets and tubes to isolate compartments hosting organs and tissues relying on strong and specific cell-cell contacts. These cells are polarized resulting in the creation of distinct apical, basal, and lateral membrane domains, each with unique molecular and functional characteristics.^8^ Apico-basal polarity of the cells dictates the placement of adhesion molecules to connect cells laterally and the arrangement of occluding junctions to prevent paracellular diffusion,^9^ forming selectively permeable barriers. Here, we show that the force response and energy dissipation depends on the cells’ polarity. When indenters are applied to the apical side of the monolayer, they initially compress the soft caps within the contact zone. Subsequently, the entire monolayer exhibits a response analogous to that observed when the basal side is targeted. Compressing the apical caps results in substantial energy dissipation (Fig. 2B), while the basal side react perfectly elastic but is slightly stiffer. At elevated force (indentation depth) the tension of the whole monolayer becomes independent of the orientation of the cell monolayer. The origin of the tension measured here is an interplay of cell-substrate anchoring, cell-cell interactions and intracellular contractility. The primary functions of cell-cell contact and cell-substrate adhesion sites are to regulate the adhesive strength. At these contact points, adhesion energy is counteracted by cell tension, predominantly mediated by the contractile actomyosin cell cortex attached to the contact.^25^ This interplay between adhesion energy and cortical tension forms the interfacial energy at the contact. If the cell-cell interaction becomes stronger the contact area expands. The same is true for the cell-substrate interaction. Therefore, adhesion energy and cortical tension are crucial parameters in determining how cells interact with their environment.^25^ Hence, it is important to notice that for the aperture-spanning monolayer, a component of the measure tension *σ*_0_ arises due to the differential free energy of surface adhesion between the aperture rims *G*_ap_ and the free-standing part *G*_cell_ of the monolayer:

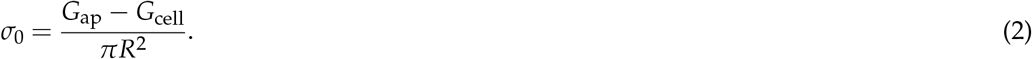

Therefore, the measured tension *σ* partly reflects this free energy difference.

Harris *et al*. were the first to examine tissue mechanics using cell monolayers spanned between two rods serving as force transducers.^4^ The authors discovered that cells construct a tissue with a significantly elevated elastic modulus (*≈*20 kPa) compared to individual cells (*≈*1 kPa).^4, 26^ The device was used to illustrate that monolayers can endure remarkably extensive deformations, frequently accommodating several-fold increases in length before rupture occurs.^6, 7^ Under significant deformation, epithelial tissues enhance their stiffness through a process regulated by a supracellular network of keratin filaments.^7^

Our experimental conditions were not suitable to advance into a regime where we could probe the stability of tissue or even obtain its 2D elastic modulus. However, we could precisely measure the lateral tension originating from differential adhesion and an intricate interplay of connectivity and contractility. Interestingly, despite being contractile, epithelial cells form monolayers that exhibit extensile behavior. This means the net force from neighboring cells and substrate interactions acts to elongate the cell further along its long axis.^27^ It was found that strong cell-cell connections are responsible for this behavior. Altering the balance between contractile and extensile forces with blebbistatin or calyculin leads to changes in tissue tension, either reducing or increasing it (Fig. 2F). Calyculin A, a serine/threonine protein phosphatase inhibitor, is known to enhance the intrinsic tension of a free-standing monolayer to the point of causing local ruptures at cell-cell junctions, a phenomenon we also observed.^7^ Conversely, blebbistatin, a myosin inhibitor specifically for myosin II, reduced tension compared to untreated cells. Recently, Fischer-Friedrich and coworkers reported that basement membrane of the wing disk epithelium of Drosophila plays a pivotal role in generating basal tension. Using atomic force microscopy, they also found that the basement membrane functions as a solid-like sheet undergoing expansile stretch to maintain epithelial homeostasis.^15^

In conclusion, our findings demonstrate that apico-basal polarity significantly influences tissue mechanics. The compliant apical side dissipates energy to mitigate stress, while the basal side, being entirely elastic, restores the tissue’s shape after dilation.

## Material and Methods

### Cell Culture

Madin-Darby Canine Kidney II (MDCK II) cells were cultivated in Minimum Essential Medium (MEM; Gibco, MT, USA) with EarleâĂŹs Salts, 2 mM GlutaMAX (ThermoFisher Scientific, MA, USA) and 10% fetal calf serum (FCS; BioWest, NuaillÃl’, France) at 37^*°*^C and 5% CO_2_ atmosphere. Subcultivation was done every two days depending on the confluency (*∼* 90%): The cells were washed with phosphate-buffered saline (PBS; Biochrom, Berlin, Germany) without Ca^2+^ and Mg^2+^ and incubated with 2 mL trypsin/EDTA (0.25%/0.02% (*w*/*v*); BioWest/Biochrom) solution at 37*°*C for 5 min. The reaction was stopped by the addition of 1 mL culture medium and 1 mL FCS. Detached cells were centrifuged at 270 *×g* for 3 min, the supernatant was aspirated, and the pellet resuspended in 1 mL culture medium. Afterwards the cells were counted using Countstar Rigel S5 (ALIT Life Science Co. Ltd., Shanghai, China).

### Device Fabrication

Separate mounts were used for initial proof of concept to host free-standing cell monolayers (Fig. 1, Fig SI1/2). The small scale testing device adapted from Harris *et al*.^4^ was used for imaging due to its lower height, while the larger grid mount was used for force spectroscopy.

#### Small Scale Testing Device

The design consist on a block with two glass rods attached, in a higher level than the bottom (on the basis as published by Harris *et al*^4^) as shown in Fig. SI1. 3D-printed polylactic acid (PLA; Ultimaker, Utrecht, Netherlands) blocks with two inserted borosilicate glass rods (GB100F-8P; Science Products GmbH, Hofheim, Germany) were used. The blocks, designed with FreeCAD version 0.21, have the dimensions of 7 *×* 5 *×* 4 mm^3^ (L *×*D *×*W), while the recesses of the holes have a depth of 3 mm and a distance of 1.3 mm. The rods were cut to a length of approximately 1 cm and fixated in two holes with two-component glue (eco-sil; Picodent, Wippenfürth, Germany), before the whole mount was glued to an culture dish (*µ*-Dish 35 mm low glass; ibidi, Martinsried, Germany). After curing, the dishes are sterilized for 15 min with 2 mL of 70% ethanol p.a., washed two times with 2 mL sterile water and dried for at least 30 min.

#### Grid Device for Atomic Force Microscopy

The second design involves a stainless steel tripod made to fit in the inner ring of an ibidi culture dish (*µ*-Dish 35 mm high glass; ibidi) (Fig. 1 and Fig. SI2). Atop of the small supporting surface of the tripod rests a stainless steel grid, topped with a borosilicate glass ring. The grid functions as a support for the collagen matrix, while the glass ring confines the area, which is filled up with collagen and offers an adhesive surface for cell attachment. The design allows to remove the grid after digestion and to flip the glass ring with the attached monolayer, which can be placed back into the tripod. The cellular layer remains attached to the bottom of the glass ring, and it provides also access to the basal side. Stainless steel parts were autoclaved, while the glass ring was sterilized with UV light (*λ* 365 nm) for 1h. The parts were assembled under sterile conditions in an ibidi high glass culture dish.

### Cell Culture on Mounts

Cellmatrix^®^ collagen type I-A (Nitta Gelatin Inc, Osaka, Japan) was reconstituted on ice with cooled pipette tips to a final concentration of 1.5 mg mL^-1^. For this, collagen, ddH_2_O, 10 *×*MEM and sterile reconstitution buffer (2.2 g NaHCO_3_ in 100 mL of 0.02 N NaOH and 200 mM Hepes) were mixed thoroughly in a ratio of 5:2:2:1.^4^ The collagen matrix is prepared freshly for each experiment.

For the testing mount, 12 *µ*L reconstituted collagen is deposited between the glass rods as described by Harris *et al*.^28^ For the grid mount, 200 *µ*L of collagen matrix were placed onto the grid with a 200-*µ*L pipette and cooled tips. The matrix was stroked carefully with the side of a cooled pipette tip up to the glass ring, by slightly tilting the ibidi culture dish. Here, it is ensured that the matrix reaches up to the glass ring, but collagen spreading on top of the glass is avoided. Afterwards, the collagen is allowed to dry for 45-60 min at 37*°*C, 5% CO_2_.

Then 29,000 MDCK II cells in 4 *µ*L culture medium with 0.1% Penicillin/Streptomycin solution (Pen/Strep; Merck, Darmstadt, Germany) are prepared and seeded onto the collagen support of the testing mount, while for the grid mount 3 *·* 10^5^ cells in 180 *µ*L were used, respectively. Furthermore, for both mounts 10^5^ cells in 300 *µ*L culture medium with 0.1% Pen/Strep solution were seeded in droplets to the outer ring of the ibidi culture dish. The mounts were incubated for 1h at 37*°*C, 5% CO_2_ to achieve attachment of the cells. Subsequently, the different ibidi culture dishes were filled up with culture medium with 0.1% Pen/Strep solution until the mounts were completely submerged (2 mL for the testing mount in ibidi low glass bottom and 4 mL for grid mount in ibidi high glass bottom *µ*-dishes).

After 48h incubation of the testing mount and 72h incubation of the grid mount, the cellular layer was washed once with culture medium before the collagen matrix was removed by enzymatic digestion with 1 mg mL^-1^ collagenase type II (Gibco, MT, USA) solution in culture medium. This was followed by an incubation of 40âĂŞ60 min at 37*°*C and 5% CO_2_, during which the success of the digestion is monitored visually. The collagenase solution is then partially replaced by culture medium and the cell layer incubated for another 10âĂŞ20 min. Afterwards, a tripod in an ibidi high glass culture dish equipped with 4 mL of culture medium with 15 mM Hepes and 100 *µ*L Pen/Strep solution was prepared. The glass ring with the attached monolayer was carefully and quickly transferred with two tweezers to the new culture dish mounted on the tripod.

#### Immunostaining

The cells are fixed with paraformaldehyde (4% in PBS; Science Services, Munich, Germany) for 20 min at room temperature and permeabilized with Triton X-100 (0.1% in PBS; Sigma Aldrich, MO, USA) for 5 min at room temperature. A blocking solution consisting of 2% (*w*/*v*) BSA (Carl Roth GmbH + Co. KG, Karlsruhe, Germany) with 0.1% Tween 20 (Sigma Aldrich, MO, USA) (*v*/*v*) is applied for 2h at room temperature. To proof the efficiency of the collagenase digest, a cellular layer of MDCK II with GFP-labeled Myosin (kind gift of Prof. Alf O. Honigmann, Technical University of Dresden, Germany) on the smaller testing mount is cultivated and the collagen matrix stained with anti-collagen type I antibody (clone 5D8-G9, 1:100, 1h; Merck Millipore, MA, USA) without its supporting matrix. As secondary antibody goat anti-mouse IgG AlexaFluor546 is used (1:400, 1h; Life Technologies, CA, USA). To visualize the polarity, the free-standing monolayer grown on the small testing mount is immunostained after the collagenase treatment with anti-podocalyxin (1:100, 1h; Sigma Aldrich, MO, USA) and anti-laminin (1:100; Sigma Aldrich, MO, USA) antibodies. As secondary antibodies, goat anti-mouse IgG AlexaFluor546 or goat anti-rabbit IgG AlexaFluor488 (both 1:400, 1h; Life Technologies, CA, USA) are used, respectively. DAPI diluted 1:10 (c_stock_ 500 ng mL^-1^) (Sigma Aldrich, MO, USA) in methanol for 15 min stains the nuclei.

### Drug Treatment

Different drugs acting on the cytoskeleton are applied to the free-standing monolayer 15 min before the force measurements. For this, 2 mL of culture medium are removed from the ibidi culture dish directly at the AFM and replaced by 2 mL of the respective compound dissolved in culture medium with 15 mM Hepes and 100 *µ*L Pen/Strep solution. To induce a cell softening and a reduction in cortical tension, the inhibitor of class II myosins para-aminoblebbistatin (Bleb; 50 *µ*M Motorpharma Ltd., Budapest, Hungary) was applied. Contrary, the drug calyculin A (Caly; 20 nM, Enzo Life Sciences Inc., NY, USA) inhibits the myosin light chain phosphatase, which increases the throughput of active myosin II, leading to an increase in monolayer tension and decreased fluidity, suggestive of stiffening of cells. Lastly, the organic compound glutar-dialdehyde (GDA; 0.5%, Merck KGaA, Darmstadt, Germany) was applied to the cellular layer, which crosslinks primary amino groups leading to increased rigidity.

### Atomic Force Microscopy

After removal of the collagen matrix, force spectroscopy measurements are conducted using a JPK CellHesion^®^ 300 (AFM; Bruker Nano, Berlin, Germany) mounted onto an inverted light microscope IX83 (Olympus, Tokio, Japan). All measurements were carried out at 37*°*C maintained by a Petri dish heater (Bruker Nano, JPK). The used cantilever have a spherical tip of borosilicate glass with a diameter 20 *µ*m and and a spring constant of 0.5-9.5 N m^-1^ (CP-FM-BSG-C; sQUBE, Bickenbach, Germany). Force curves were obtained at a speed of 2 µm s^-1^ and a sample rate of 4000 Hz at constant height mode with dynamic baseline adjustment. The approach to the surface was done manually before force-distance and force-time curves for different setpoints (30-200 nN) were recorded. To ensure that no time dependencies influence our results, the measurements were started alternately on the apical and basal side. To switch to the opposite domain, the glass ring is removed from the medium and placed in a new culture dish with a tripod and culture medium with 15 mM Hepes and 100 µL Pen/Strep solution as described above. The total measurement time was limited to 3h to ensure a maintenance of polarity.

### Model for Indentation of a free-standing Monolayer

Here, we describe how cell monolayers spanned over a circular aperture respond to indentation with a spherical indenter leading to the following contour Fig. 2A. *R* denotes the radius of the aperture and *a* is the contact radius of the monolayer with the conical indenter.

#### Free Energy of a free-standing Cell Monolayer

The free energy of the cell monolayer treated as a continuous sheet of a homogeneous material is given by:^29, 30^

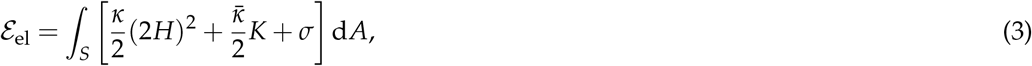

where *S* is the surface of the membrane, *H* its mean curvature (2*H* = 1/*R*_1_ + 1/*R*_2_) and *K* the Gaussian curvature (*K* = (*R*_1_*R*_2_)^*−*1^). *κ* and 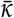 are the splay and saddle splay moduli, respectively. For the sake of simplicity, we neglect the Gaussian modulus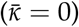.^31^ *σ* denotes the tissue tension comprising mainly the free energy contribution (per unit area) arising from adhesion of the cells to the pore rim, and in more general terms it represents the chemical potential of the monolayer reservoir recruited during deformation. It can also be considered the Lagrangian multiplier to keep the area constant. The shape equation is obtained from standard variational calculus representing the balance of normal forces per unit area:^32^

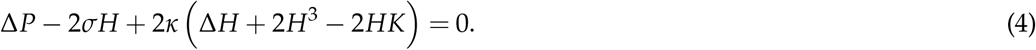

The pressure difference Δ*P* enters as the Lagrange multiplier ensuring constant volume. For free-spanning monolayers on pores open to both sides we can discard this contribution. We assume that the energy contribution due to recruiting new surface area against the tissue tension is substantially larger than the bending energy and we can rewrite equation (4):^31^

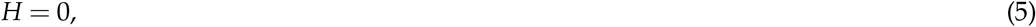

the minimal surface equation. The problem of finding the shape *r*(*z*) of the membrane during indentation therefore reduces to the problem of finding its minimal free surface.^31, 33^ We model the contact zone of the spherical indenter with the cell monolayer as a perfect ring. We consider the elementary case of two equally sized rings separating the monolayer by 2*L* forming a shape with zero mean curvature. The surface area is

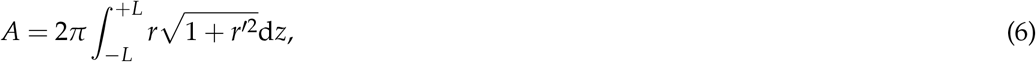

with *r*^*′*^ = d*r*/d*z*. Applying standard variational calculus we obtain:^33^

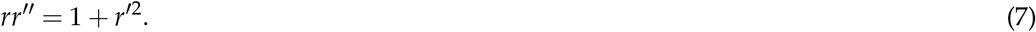

This differential equation can be integrated in two steps.^33^ Using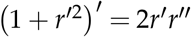 we get

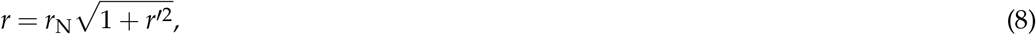

with *r*_N_ being a constant that contains the restoring force (*vide infra*). Second, we employ the identity

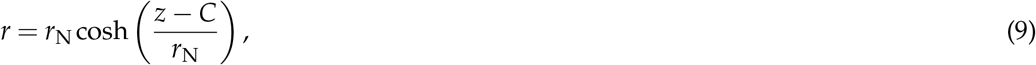

with the integration constant *C* being zero for two equally sized rings.^33^ Since we have one radius given by the aperture *R* and the other one by the contact radius *a* with the indenter, we need to calculate *C* from this boundary condition. *r*_N_ is identified as the minimal radius of the catenoid, its so-called neck radius. *C* can be obtained from *r* = *R* of the upper ring

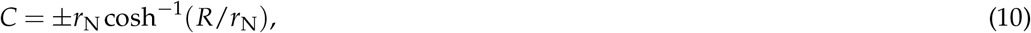

leading to

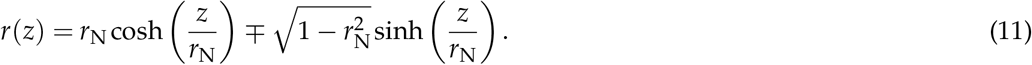

The solution is a catenoid with two differently sized rings in which the upper sign corresponds to catenaries with a minimum neck radius at a positive value of the indenter depth *z*(*C >* 0). For the shape of the free-standing monolayer (*z*(*r*)) we find

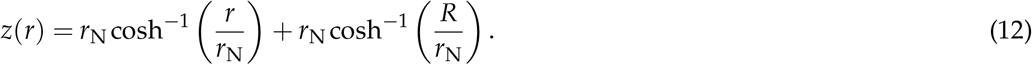

Writing it in nondimensional form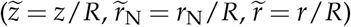 we obtain:

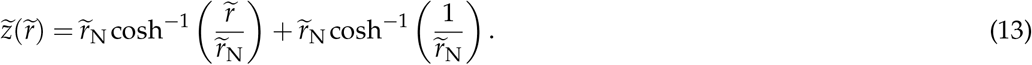

The relation between *r*_N_ the minimal radius and the force *f* reads

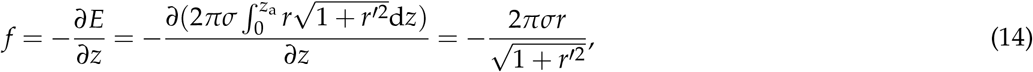

with *z*_*a*_ the indentation depth at the contact line *r* = *a*. At the neck of the catenoid *r* = *r*_N_, we have *r*^*′*^ = 0 and therefore *r*_N_ = *f* /(2*πσ*). Since at *r*(*z*_*a*_) = *a* with *a > r*_N_ we can write:

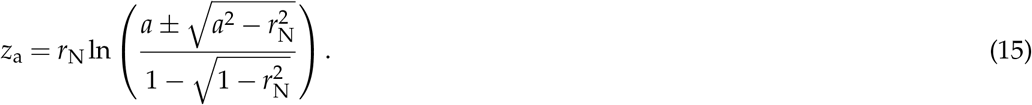

Equation 15 is responsible for two branches forming a closed curve in the *f − z*_a_ plane for *a < R*.^31^ A critical (maximal) separation *z*_a_ exists, where no solution exists and the catenoid is unstable. If the indentation depth is below this critical separation two catenoidal equilibrium solutions can be found (see equation 15, see Fig. 2C). We only consider the branch with larger *r*_N_ that has lesser area (minus sign in equation 15). Now we are left to find the contact radius *a* from the continuity condition, where the slope is identical for indenter and free-standing membrane. For a spherical indenter we find

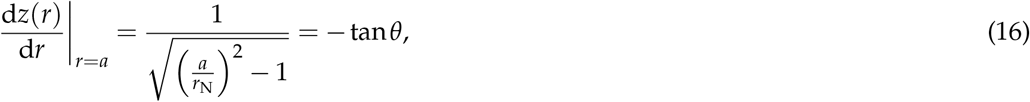

with

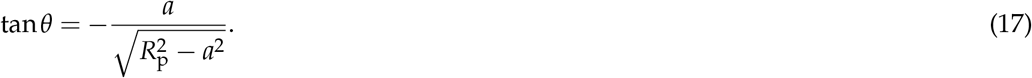

Here, *R*_p_ is the radius of the indenter. We now have to solve

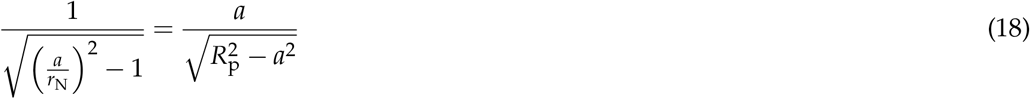

numerically to obtain *a*. The indentation depth at the tip of the indenter is *z*(*r* = 0) = *z*_a_ + *a* tan(*θ*). Note, it is required that *a > r*_N_ since *r*_N_ is the smallest possible radius of the catenoid (see Fig. 2C). In non-dimensional form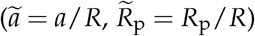 the indentation depth at *r* = 0 is

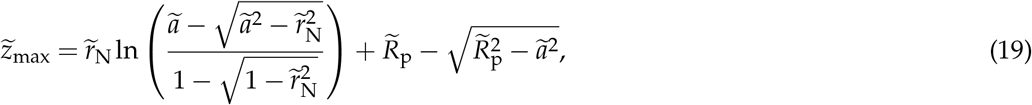

which tells us that for a given indenter geometry the shape of the membrane and its scaled force response is uniquely defined by the distance between the two rings. This allows us to first compute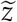 for each value of *R* and subsequently fit the two curves with a polynomial,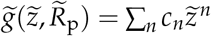to obtain an analytical solution of the elastic response to indentation. For the approach force curve we get (negative sign introduced to capture the experimental data,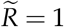):

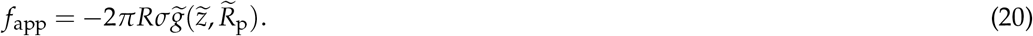

In the limit of small forces we obtain:

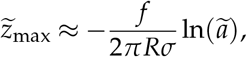

If we further approximate the contact radius with the radius of the indenter (*a*_max_ = *R*_p_) we get:

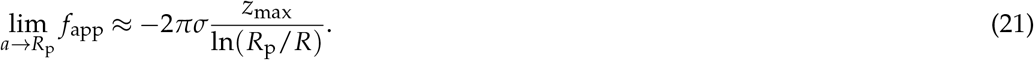

### Slack of the Monolayer

It is found experimentally that the planar, free-standing cell monolayer is subject to gravitation and shows a slack on the order of 100 *µ*m (see Fig. 3B). The homogeneous force per area of the aperture (*P* = *F*/*A*) acting on the monolayer is its own weight, i.e., gravity (*F* = *mg* = *ρ*Δ*A*Δ*zg*) :

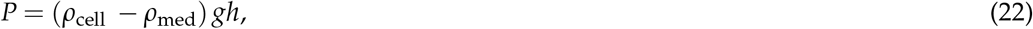

with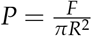·*h* is the thickness of the cell layer, *ρ*_cell_ the density of the cell, *ρ*_med_ the density of the medium, accounting for the buoyancy of the cell-monolayer), and the constant of gravity *g* = 9.81 m/s^2^. *z*(*r*) is the deflection of the cell layer at any radial position *r*. The corresponding Euler-Lagrange equation to be solved to describe the shape of the cell layer is:

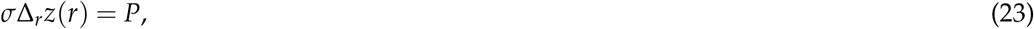

with *σ*, the tissue tension and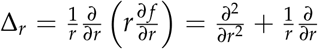, the Laplace operator in circular coordinates assuming axisymmetry. This gives:

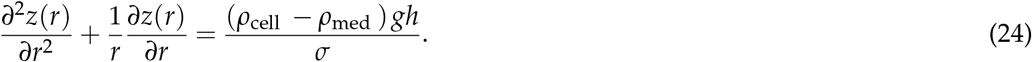

The boundary condition at the rim is:

We can solve this as:

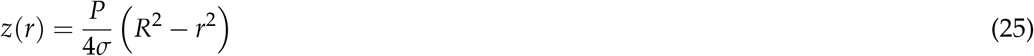

Hence the maximal slack (*z*(*r* = 0) = *z*_max_) is

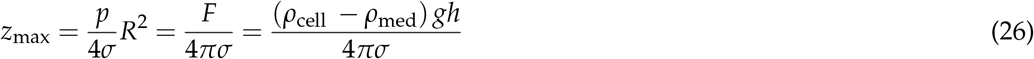

Estimation of the gravitational force for the given geometry of the described experiment:

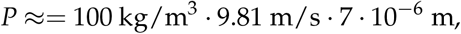

which is

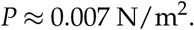

The force acting on an aperture with *R* = 10^*−*2^ m is:

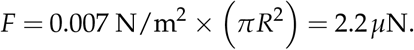

Assuming *σ* = 2 mN/m, we obtain for the maximal slack:

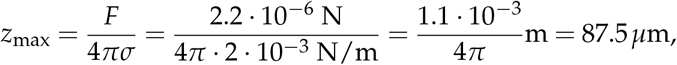

which is in good accordance with experimental findings.

#### Calculation of the Tension

To evaluate the force-height curves, a self-written Python script was used to correct for the baseline and determine the contact point by using Knee algorithm.^34^. The curves are extracted from the contact point to the maximum force (near the selected setpoint). The slope *k* is then taken from the 50% to the 90% of these curve, where a linear fit is applied. The tension *σ* becomes then

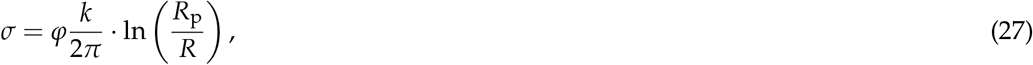

where *R*_p_ corresponds to the radius of the indenter and *R* to the radius of the complete monolayer (inner ring size). *φ* = 1.15 is a correction factor empirically determined by using the full theory (equation (20), see Fig. SI6) to generate force versus indentation data fitted by equation (21).

### Statistical Analyses

Values and error bars are displayed as mean *±* standard deviation. The data were evaluated for their differences using the Mann-Whitney Wilcoxon rank sum test (ns: *p*-value *>*0.05, * denotes *p <*0.05, ** refers to *p <*0.01, *** indicates *p <*0.001, and **** imply *p <*0.0001).

## Conclusion

The response of planar free-standing cell monolayers to central deformation with a colloidal probe is examined as a function of apico-basal polarity and intrinsic tension, using cytoskeletal drugs to interfere with actomyosin contractility. Cells were found to be apically softer than basally and dissipate energy during indentation-retraction cycles. This mechanical polarity helps cells mitigate rising Laplace pressure in the renal tube by dissipating energy when the apical cap is compressed and recruiting excess area stored in the apical cap (e.g. in microvilli). Conversely, the predominantly elastic basal side ultimately resists deformation like an elastic, pre-stressed thin plate, providing the necessary elasticity for the tissue to restore its shape after experiencing dilation. Ultimately, the integrity of the tissue is conserved by cell-cell connections, which were not probed by this relatively mild deformation.

## Acknowledgments

The authors thank Dr. Tabea A. Oswald and Angela Rübeling for their help with cell culture.

## Author contributions statement

A.P-T. and A.J. conceptualized the idea, A.P-T. and U.U. performed the colloidal probe measurements, A.P-T. analyzed the data. A.J. wrote the manuscript and developed the model. All authors contributed in editing and writing of the manuscript.

## Supplementary Information

### 1 Design of the Small Device

**Figure 1:**
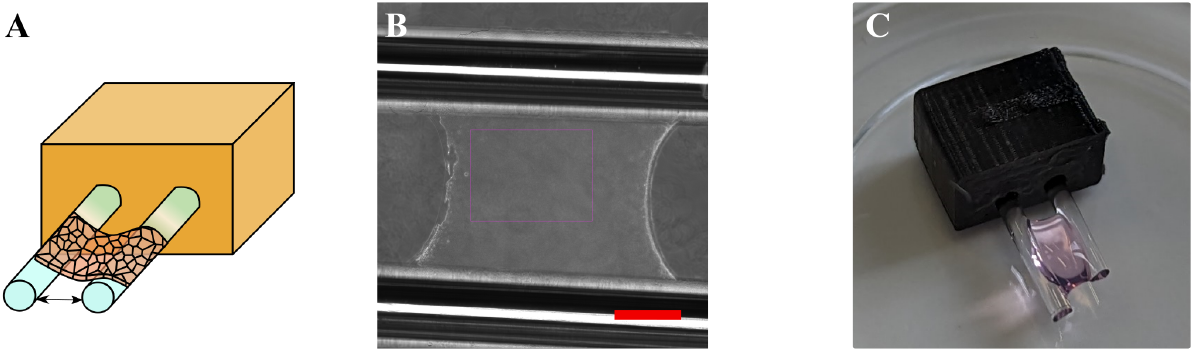
Fabrication of the small testing device. A) Scheme of the PLA (polylactic-acid) block containing two glass rods. B) Phase contrast image of MDCK II cells attached to the glass rods, forming a planar tissue. Scale bar: 1 mm. C) Picture of the device and the free-standing collagen matrix.

### 2 Design of the Grid Device

**Figure 2:**
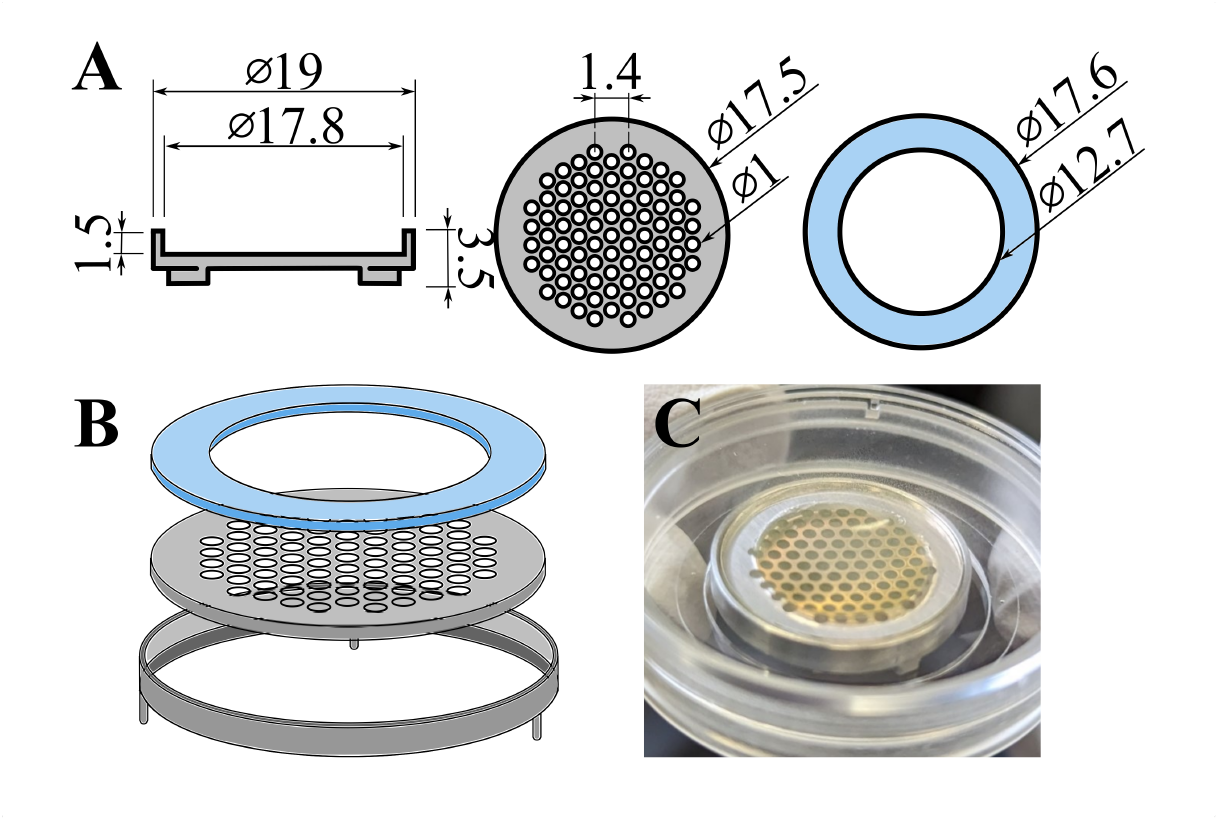
Scheme of the grid device construction to host free-standing cell monolayers. A) The mount includes a tripod made of stainless steel, designed to fit in an ibidi Petri dish. B) A small support surface carries a stainless steel grid and on top a glass ring. The grid serves as a support for the collagen matrix. The glass ring (height: 1 mm) confines the area, filled up with collagen and provides space for attachment of cells. C) Exemplary image of the mount filled with collagen is shown.

### 3 Indentation on Pure Collagen Gel Surface and Collagen with Cells

**Figure 3:**
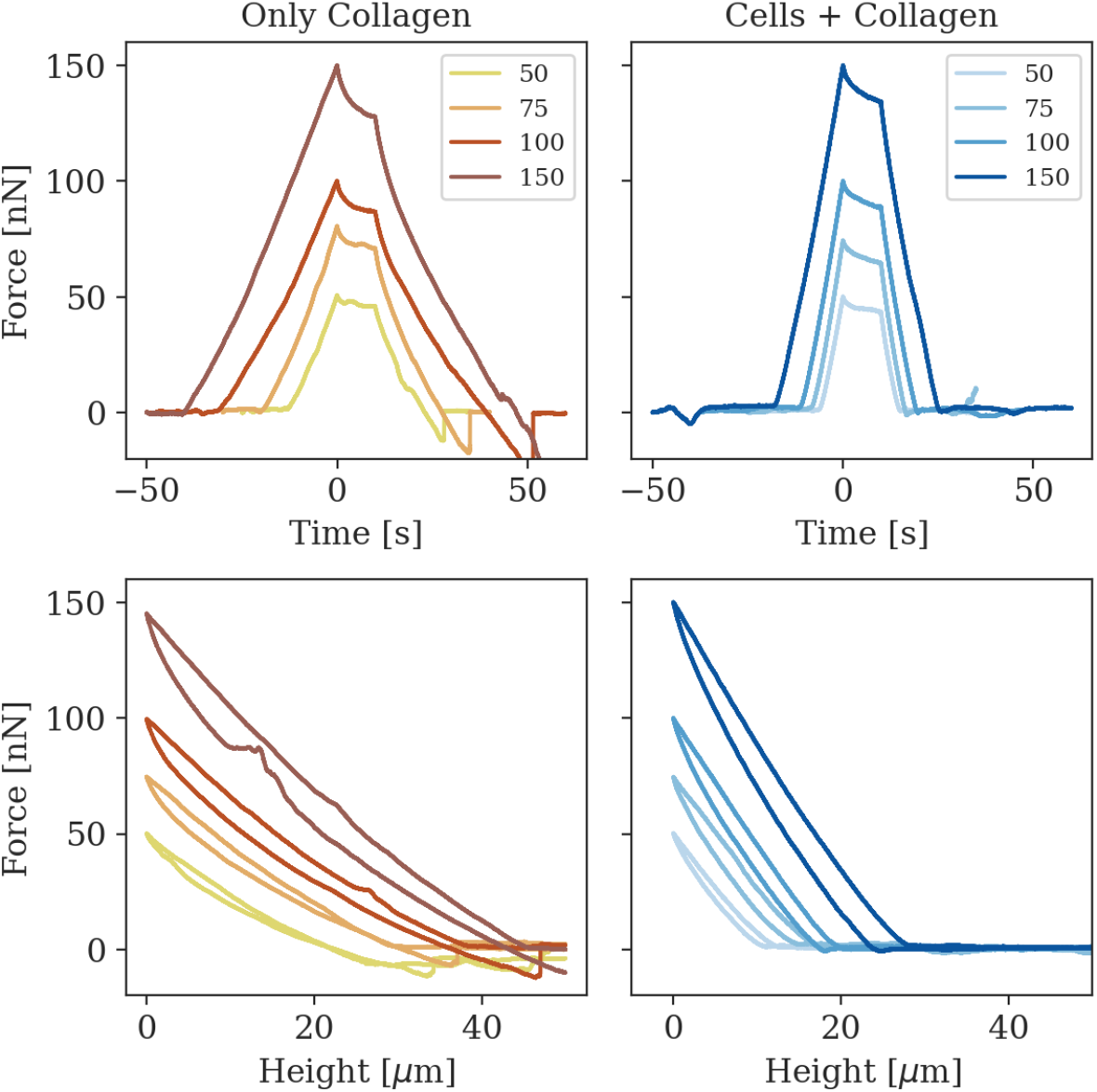
Exemplary force-time curves (upper row) of indentation-relaxation experiments and their homologous force-height curves in indentation-retraction (bottom row) from colloidal probe force measurements on the collagen matrix (left) and a cellular layer on top of the collagen matrix (right) using different maximal force setpoints (50, 75, 100 and 150 nN).

**Figure 4:**
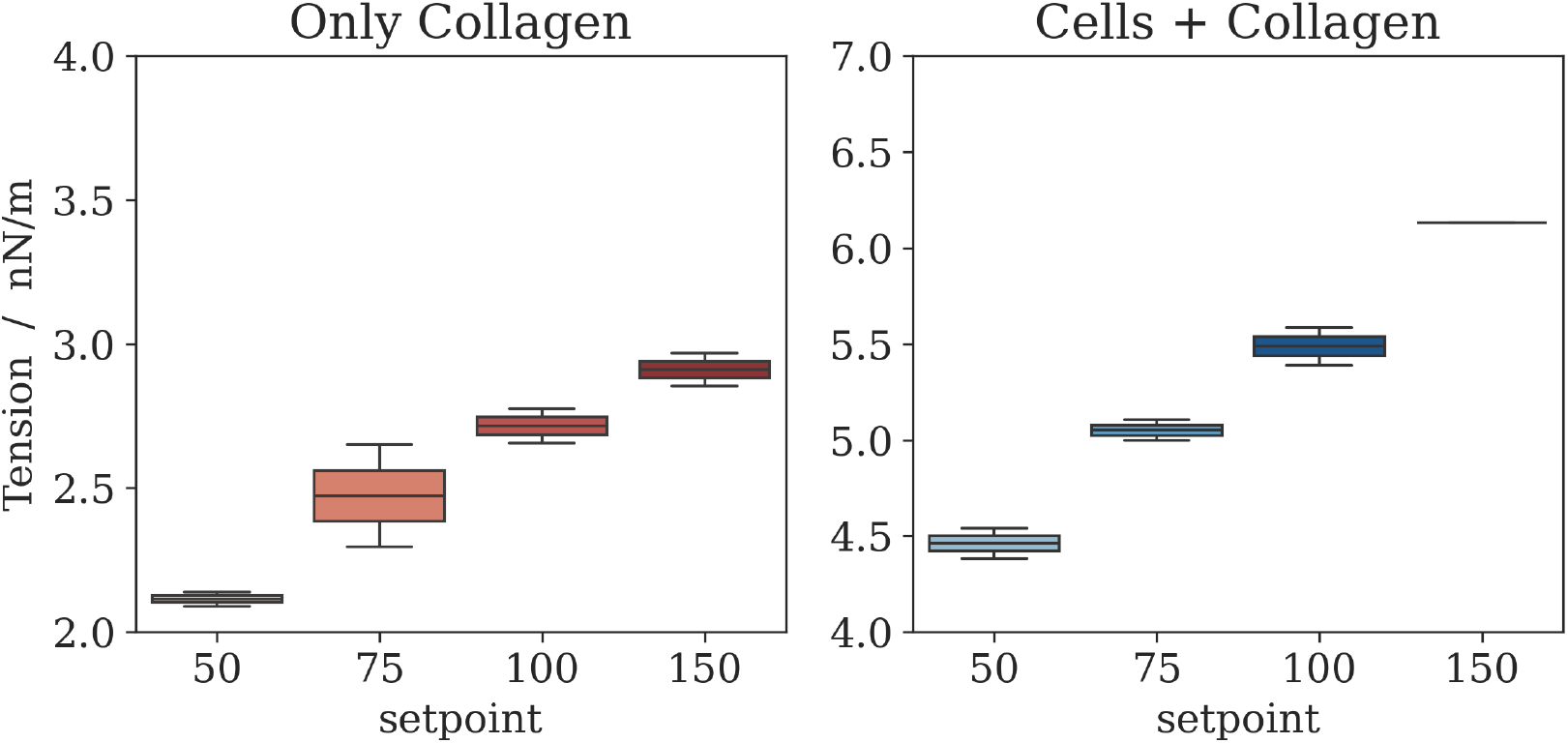
Tension of force-time curves recorded on the collagen matrix and a cellular layer on top of the collagen matrix for different maximal force setpoints (50, 75, 100 and 150 nN). In the diagram the line within the box is the median value for each case.

### 4 Hysteresis only appears at Low Indentation Depth

**Figure 5:**
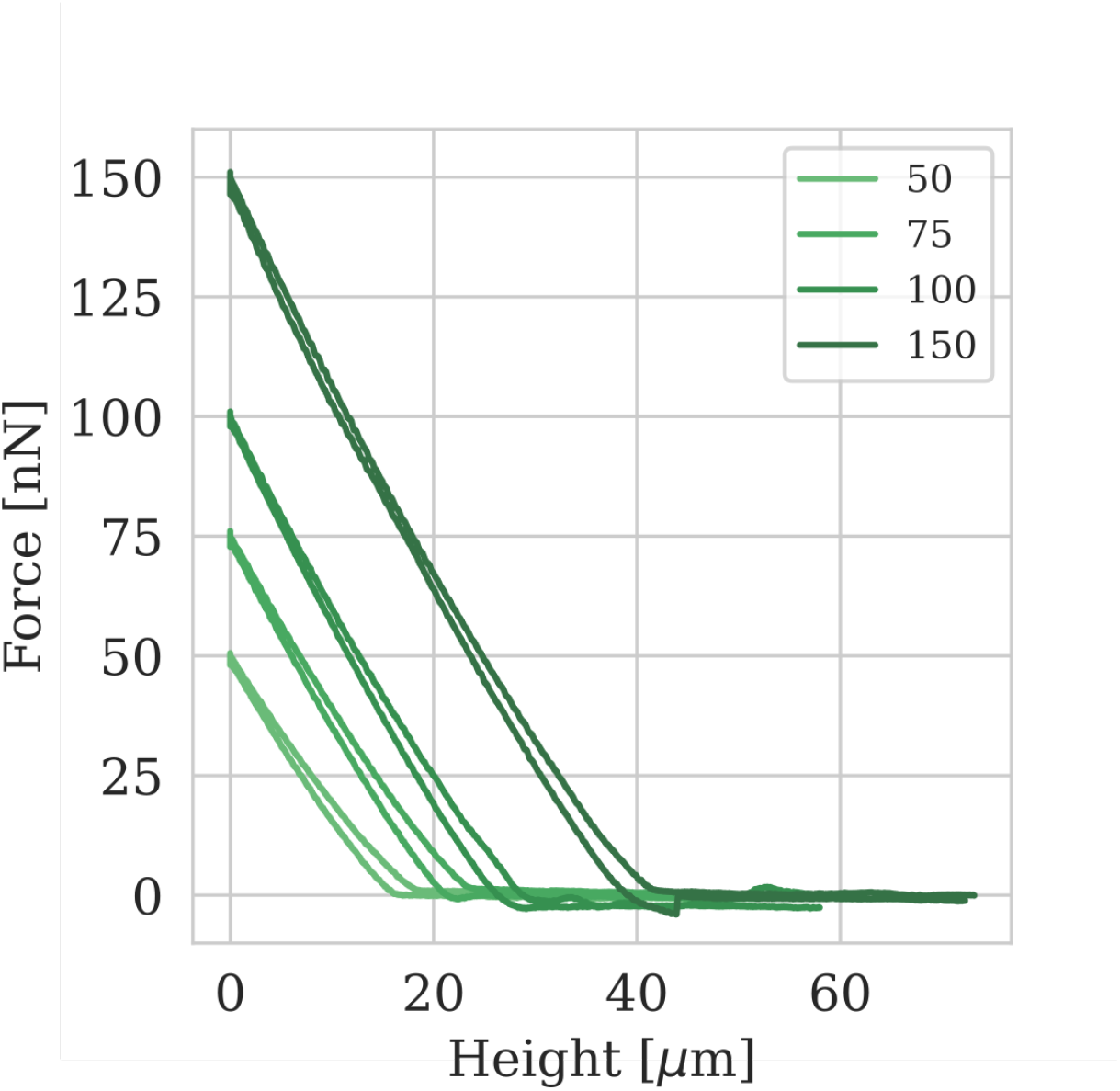
Force indentation curves obtained from the apical domain of a free-standing cell monolayer reaching different maximal indentation depths (height). This was reached by using different maximal force setpoints (50, 75, 100 and 150 nN). Notably, at large indentation depths the hysteresis vanishes and the response becomes elastic.

### 5 Correction Factor for obtaining Tissue Tension

Tissue tension is obtained from energy minimization resulting in a minimal surface of the free part of the cell monolayer. The resulting force-indentation relation in the limit of small forces reads

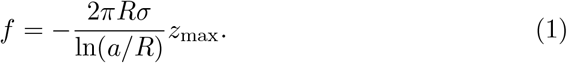

If we further approximate the contact radius with the radius of the indenter *a*_max_ = *R*_p_, (note that the contact radius cannot become larger than *R*_p_) we get:

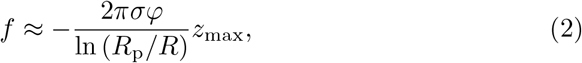

with *φ*, an empirical correction factor obtained from fitting eq. (2) to data obtained from evaluating eq. (1).

**Figure 6:**
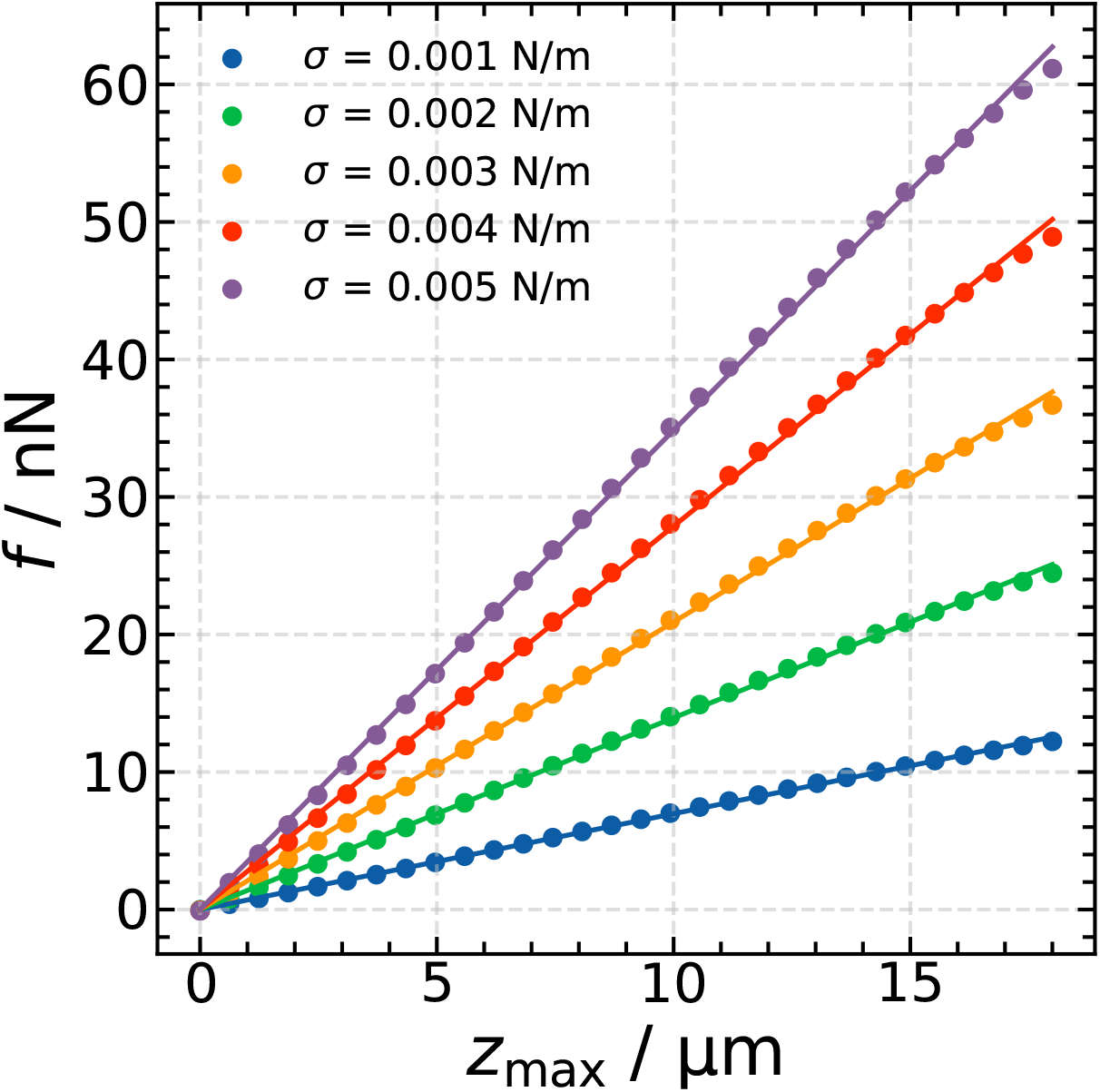
Linear fit of data from the full solution *f* (*z*_max_) (eq. (1), dots) for various values of *σ* to obtain the correction factor *φ*. Here, in this range of indentation depth and tissue tension, we find *φ* = 1.15.

